# Genome-wide association of PCSK9 (Proprotein Convertase Subtilisin/Kexin Type 9) plasma levels in the ELSA-Brasil study

**DOI:** 10.1101/2020.12.03.409631

**Authors:** Isabela Bensenor, Kallyandra Padilha, Isabella Ramos Lima, Raul Dias Santos, Gilles Lambert, Stéphane Ramin-Mangata, Marcio S Bittencourt, Alessandra C Goulart, Itamar S. Santos, Jose G Mill, Jose E Krieger, Paulo A. Lotufo, Alexandre C. Pereira

## Abstract

Pharmacological inhibition of PCSK9 (proprotein convertase subtilisin/kexin type 9) is an established therapeutic option to treat hypercholesterolemia and plasma PCSK9 levels have been implicated in cardiovascular disease incidence. A number of genetic variants within the PCSK9 gene locus have been shown to modulate PCSK9 levels, but these only explain a very small percentage of the overall PCSK9 interindividual variation. Here we present data on the genetic association structure between PCSK9 levels and genome-wide genetic variation in a healthy sample from the general population.

We performed a genome-wide association study of plasma PCSK9 levels in a sample of Brazilian individuals enrolled in the ELSA-Brasil cohort (n=810). Enrolled individuals were free from cardiovascular disease, diabetes and were not under lipid-lowering medication. Genome-wide genotyping was conducted using the Axiom_PMRA.r3 array and imputation used the TOPMED multi-ancestry sample panel. Total PCSK9 plasma concentrations were determined using the Quantikine SPC900 ELISA kit.

We observed two genome-wide significant loci and seven loci that reached the pre-defined *p* value threshold of 1 × 10^−6^. Significant variants were near *KCNA5* and *KCNA1*, and *LINC00353*. Genetic variation at the *PCSK9* locus was able to explain approximately 4% of the overall interindividual variation in PCSK9 levels. Colocalization analysis using eQTL data suggested *RWDD3*, *ATXN7L1*, *KCNA1*, and *FAM177A1* to be potential mediators of some of the observed associations.

Our results suggest that PCSK9 levels may be modulated by *trans* genetic variation outside of the *PCSK9* gene and this may have clinical implications. Understanding both environmental and genetic predictors of PCSK9 levels may help identifying new targets for cardiovascular disease treatment and contribute to a better assessment of the benefits of long-term PCSK9 inhibition.

## Introduction

Proprotein convertase subtilisin/kexin type 9 (PCSK9) is a key modulator of LDL receptor (LDLR) degradation and, consequently, LDL-cholesterol (LDL-C) serum levels. Gain-of-function mutations in *PCSK9* have been shown to cause familial hypercholesterolemia and increased cardiovascular risk [1]. On the other hand, loss-of-function variants have been shown to associate with low LDL-C levels and reduced cardiovascular risk [2]. Furthermore, plasma PCSK9 has been independently associated with other components of the lipid profile [3,4]. As a result, pharmacological inhibition of PCSK9 became mainstream as a lipid reduction strategy [5].

Understanding the factors that modulate interindividual variability of PCSK9 plasma levels is important for the better understanding of individual responses to treatment as well as the identification of new targets for cardiovascular disease treatment. The use of unbiased genetic approaches has the potential to contribute to increase our understanding of both.

Here we have conducted a genome-wide association study (GWAS) in healthy individuals from the general population aiming at the identification of genetic variation associated to plasma PCSK9 levels.

## Materials and methods

### Study population

The study sample belongs to the ELSA-Brasil (Estudo Longitudinal de Saude do Adulto, NCT02320461). For the present analysis we used 810 participants that have both PCSK9 plasma levels and genome-wide genotype information.

The ELSA-Brasil study design and cohort profile have been published elsewhere [6]. Briefly, ELSA-Brasil enrolled 15,105 civil servants living in six large Brazilian urban areas (Belo Horizonte, Porto Alegre, Rio de Janeiro, Salvador, Sao Paulo, and Vitoria), aged between 35 and 74 years at baseline. Information on sociodemographic, clinical history, family history of diseases, lifestyle factors, mental health, cognitive status, and occupational exposure was assessed from August 2008 to December 2010. Anthopometric, laboratory and imaging measurements were also obtained. The study was conducted in accordance with the Declaration of Helsinki and was approved by the local institutional review boards. In addition to baseline measurements samples of plasma and DNA were collected and stored for further analysis at −80 °C [7]. All participants signed an informed consent before enrollment.

Participants enrolled in the Sao Paulo site (5061 people in total) without diabetes (exclusion criteria: fasting plasma glucose-FPG > 126 mg/dL and/or 2-h post-load glucose > 200 mg/dL and/or history of treatment with oral anti-diabetic agents or insulin), without cardiovascular, renal or hepatic diseases (exclusion criteria: self-reported history of medical diagnosis of these pathologies), and who did not report prescription of lipid-lowering agents, were eligible for a PCSK9 ancillary and exploratory study. From the 1,751 randomly selected participants fulfilling the inclusion criteria for subsequent PCSK9 plasma concentrations measurements [8], we have used the 810 who had genome-wide genotype information for the present analysis.

### Biochemical analyses

A 12-h fasting blood sample was drawn in the morning soon after arrival at the research clinic, following standardized procedures for samples collection and processing. A standardized 75 g oral glucose tolerance test (OGTT) was performed in all participants without known diabetes utilizing an anhydrous glucose solution. For measurement of fasting and post-load glucose we used the hexokinase method (ADVIA 1200, Siemens); for fasting and post-load insulin, an immunoenzymatic assay; for HbA1c, high-pressure liquid chromatography. Total cholesterol (TC), high density lipoprotein-cholesterol (HDL-C) and triglycerides (TG) were measured with enzymatic colorimetric assays (ADVIA Chemistry). LDL-C was calculated using the Friedewald equation. When TG were ≥400 mg/dL, LDL-C was measured directly with an enzymatic colorimetric assay (ADVIA Chemistry)

Total PCSK9 plasma concentrations were determined using the Quantikine SPC900 ELISA kit (R&D Systems, Lille, France)^8^. Briefly, plasma samples were diluted 1:20 in the calibrator diluent onto ELISA plates and incubated for 2hrs on a plate shaker at 450 rpm. Wells were rinsed with wash buffer using an automated Hydroflex TECAN microplate washer. The detection HRP-conjugated antibody was added to each well and plates were incubated for 2 h at 450 rpm. Wells were rinsed. The TMB substrate solution was added to each well and plates were further incubated in the dark for 30 min at 450 rpm. Reactions were stopped by the addition of 0.2 N acid sulphuric solution. Absorbance was read at 450 nm with reference at 540 nm on an Infinite 200 pro TECAN plate-reader.

### Data availability

The data that support the findings of this study are available from ELSA-Brasil study on qualified request. Requests for access to more detailed summary statistics, replication results, and analytic methods will be considered by the authors.

### SNP Genotyping and Imputation

Genomic DNA extraction has been previously described [9]. ELSA-Brasil DNA samples for genotyped using Axiom_PMRA.r3 array (ThermoFisher) and genotypes annotated using the Axiom_PMRA.na35.annot.db provided at the ThermoFisher site. Genotype calling was performed using Affymetrix Power Tools. Initial VCF file contained 850483 variants fulfilled all quality criteria.

Imputation was performed using the Haplotype Reference Consortium Michigan Imputation Server using the TOPMED reference haplotype panel as reference. After imputation markers were kept if R2 > 0.8, and Minor Allele Frequency (MAF) > 0.01. A total of 11,524,071 SNPs were used for genome-wide analyses, 11,289,274 for autosomal, and 234,797 for X-chromosomal analysis.

### Colocalization analysis

For colocalization analysis we have defined a window spanning 500Kb center at the most associated variant in all regions classified as having a suggestive association signal. Information on all variants within this region was used for colocalization testing. We have used the LocusFocus (https://locusfocus.research.sickkids.ca/) analytical approach for colocalization testing. Briefly, all genes residing in each selected region with their expression quantitative trait loci (eQTL) summary statistics available in GTEx were sequentially tested for colocalization with the results obtained for PCSK9 association. As reference LD structure we used 1000 genomes 2012 European LD matrix (our sample has approximately 80% European ancestry). Colocalization was tested against all 48 tissues available in GTEx and the most significant signal was selected.

### Statistical Analysis

PCSK9 levels were log-transformed for all analyses. Baseline categorical parameters are presented using frequencies (proportions), continuous parameters are presented using mean ± SD. Before GWAS, we have adjusted a linear model for log(PCSK9) adjusting for age. The residuals of this model were used for GWAS. Confounding effects for age, sex, smoking and BMI were later tested for all genome-wide and suggestive GWA hits.

Genome-wide association analyses were conducted using plink. We have conducted two analysis one without any further adjustment and one adjusting for the first four principal components. The threshold for genome-wide significance was set to *p* <5×10^−8^. Associations with *p* <1×10^−6^ were considered as suggestive and presented as list of top SNPs.

Local association plots were created using LocusZoom [10]. Local linkage disequilibrium structure was determined using Haploview [11].

Mediation analysis was conducted for selected loci. To select markers for a genetic risk score for plasma PCSK9 levels we have determined independently associated variants at the PCSK9 genomic locus (cis-pQTL) through fitting a multiple linear regression model using 20 nominally associated markers at this loci and a stepwise variable selection procedure. Genetic risk score was derived as the sum of weighted genotypes by their final regression coefficients.

## Results

### Relationship between cardiovascular risk factors and plasma PCSK9

Clinical and laboratory characteristics of the ELSA-Brasil sample used in the present analysis are summarized in Table 1. Plasma PCSK9 levels were associated with TC (*p* = 0.0006), TG (*p* = 0.003), and LDL-C (*p* = 0.003) (Table 1).

**Table 1.**
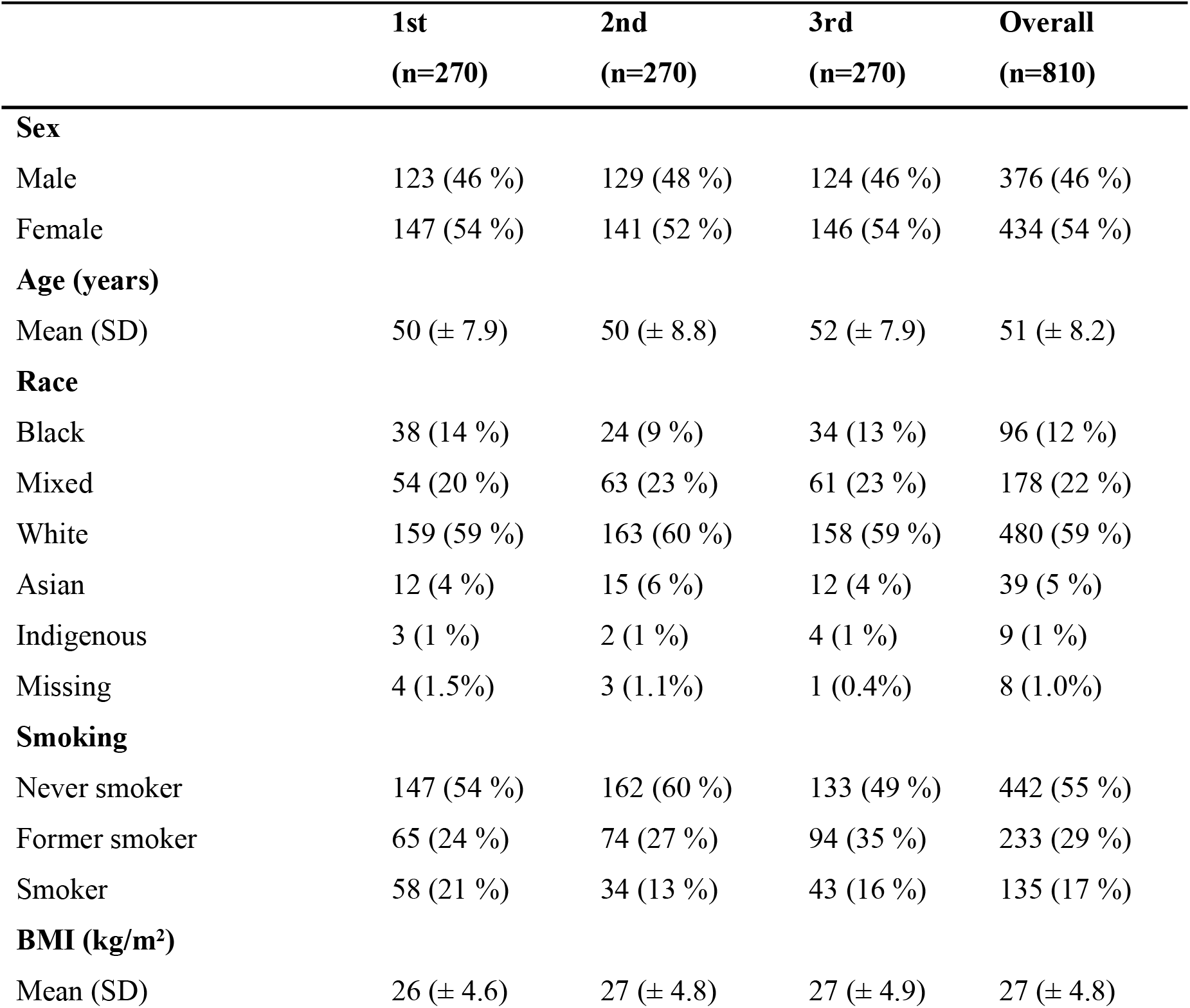
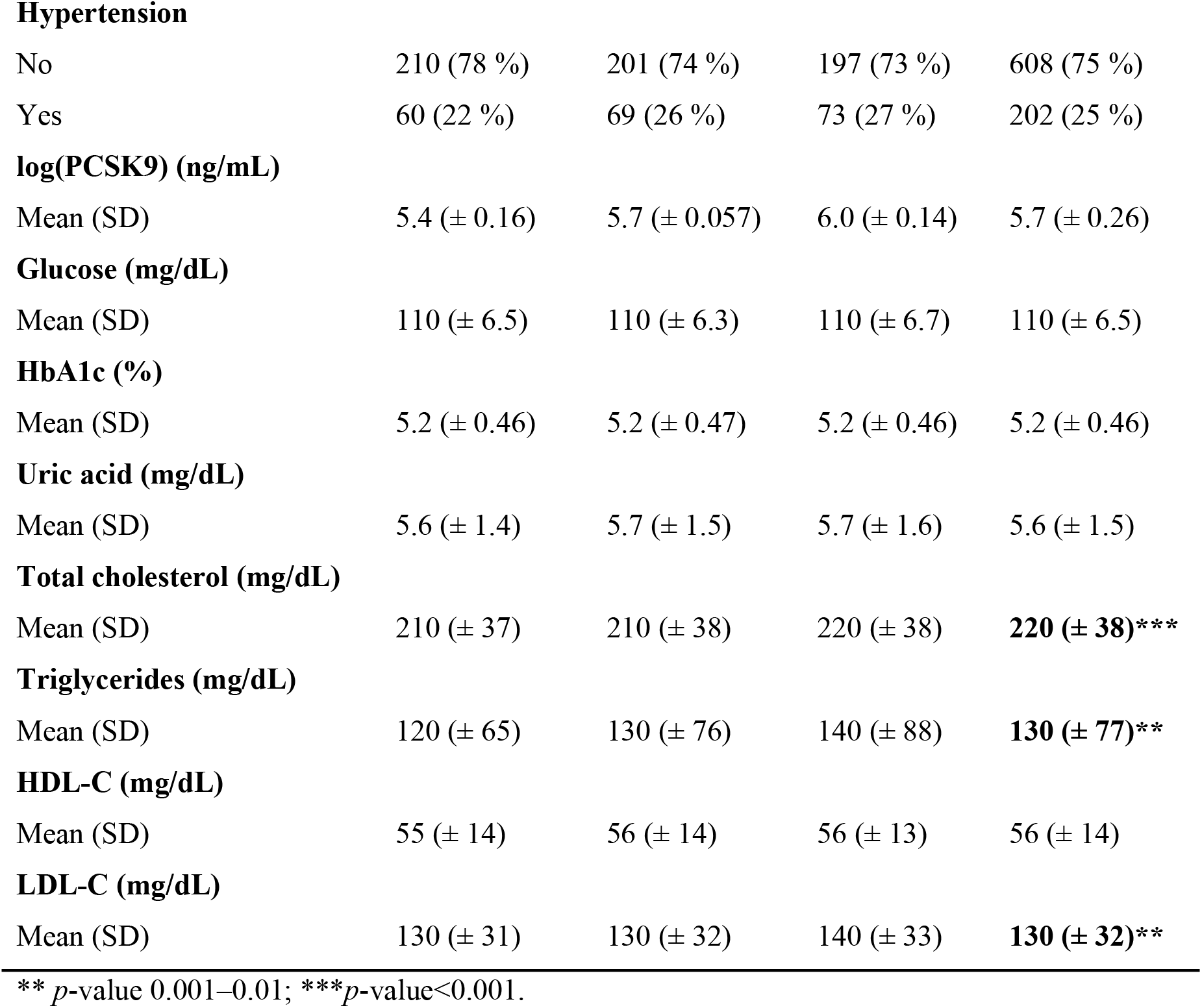
Clinical and laboratory characteristics of studied subjects according to tertiles of plasma PCSK9 concentrations.

|  | 1st<br>(n=270) | 2nd<br>(n=270) | 3rd<br>(n=270) | Overall<br>(n=810) |
| --- | --- | --- | --- | --- |
| **Sex** |  |  |  |  |
| Male | 123 (46 %) | 129 (48 %) | 124 (46 %) | 376 (46 %) |
| Female | 147 (54 %) | 141 (52 %) | 146 (54 %) | 434 (54 %) |
| **Age (years)** |  |  |  |  |
| Mean (SD) | 50 ($\pm 7.9$) | 50 ($\pm 8.8$) | 52 ($\pm 7.9$) | 51 ($\pm 8.2$) |
| **Race** |  |  |  |  |
| Black | 38 (14 %) | 24 (9 %) | 34 (13 %) | 96 (12 %) |
| Mixed | 54 (20 %) | 63 (23 %) | 61 (23 %) | 178 (22 %) |
| White | 159 (59 %) | 163 (60 %) | 158 (59 %) | 480 (59 %) |
| Asian | 12 (4 %) | 15 (6 %) | 12 (4 %) | 39 (5 %) |
| Indigenous | 3 (1 %) | 2 (1 %) | 4 (1 %) | 9 (1 %) |
| Missing | 4 (1.5%) | 3 (1.1%) | 1 (0.4%) | 8 (1.0%) |
| **Smoking** |  |  |  |  |
| Never smoker | 147 (54 %) | 162 (60 %) | 133 (49 %) | 442 (55 %) |
| Former smoker | 65 (24 %) | 74 (27 %) | 94 (35 %) | 233 (29 %) |
| Smoker | 58 (21 %) | 34 (13 %) | 43 (16 %) | 135 (17 %) |
| **BMI (kg/m<sup>2</sup>)** |  |  |  |  |
| Mean (SD) | 26 ($\pm 4.6$) | 27 ($\pm 4.8$) | 27 ($\pm 4.9$) | 27 ($\pm 4.8$) |
| **Hypertension** |  |  |  |  |
| No | 210 (78 %) | 201 (74 %) | 197 (73 %) | 608 (75 %) |
| Yes | 60 (22 %) | 69 (26 %) | 73 (27 %) | 202 (25 %) |
| **log(PCSK9) (ng/mL)** |  |  |  |  |
| Mean (SD) | 5.4 (± 0.16) | 5.7 (± 0.057) | 6.0 (± 0.14) | 5.7 (± 0.26) |
| **Glucose (mg/dL)** |  |  |  |  |
| Mean (SD) | 110 (± 6.5) | 110 (± 6.3) | 110 (± 6.7) | 110 (± 6.5) |
| **HbA1c (%)** |  |  |  |  |
| Mean (SD) | 5.2 (± 0.46) | 5.2 (± 0.47) | 5.2 (± 0.46) | 5.2 (± 0.46) |
| **Uric acid (mg/dL)** |  |  |  |  |
| Mean (SD) | 5.6 (± 1.4) | 5.7 (± 1.5) | 5.7 (± 1.6) | 5.6 (± 1.5) |
| **Total cholesterol (mg/dL)** |  |  |  |  |
| Mean (SD) | 210 (± 37) | 210 (± 38) | 220 (± 38) | **220 (± 38)\*\*\*** |
| **Triglycerides (mg/dL)** |  |  |  |  |
| Mean (SD) | 120 (± 65) | 130 (± 76) | 140 (± 88) | **130 (± 77)\*\*** |
| **HDL-C (mg/dL)** |  |  |  |  |
| Mean (SD) | 55 (± 14) | 56 (± 14) | 56 (± 13) | 56 (± 14) |
| **LDL-C (mg/dL)** |  |  |  |  |
| Mean (SD) | 130 (± 31) | 130 (± 32) | 140 (± 33) | **130 (± 32)\*\*** |

\*\* $p$ -value 0.001–0.01; \*\*\* $p$ -value<0.001.

### Genome-Wide Association Analysis of PCSK9 Plasma Levels

We have performed a GWA of the age-adjusted residuals of the log transformed values of plasma PCSK9. In the primary analysis adjusted for the four first principal components we identified two loci that reached the pre-defined genome-wide significant level of 5 × 10^−8^ (Fig 1). Notably, no significant genomic inflation was observed (lambda = 1.06). In addition, no significant difference was observed between genome-wide significant and suggestive loci when running an unadjusted analysis (S1 Fig).

**Fig 1.**
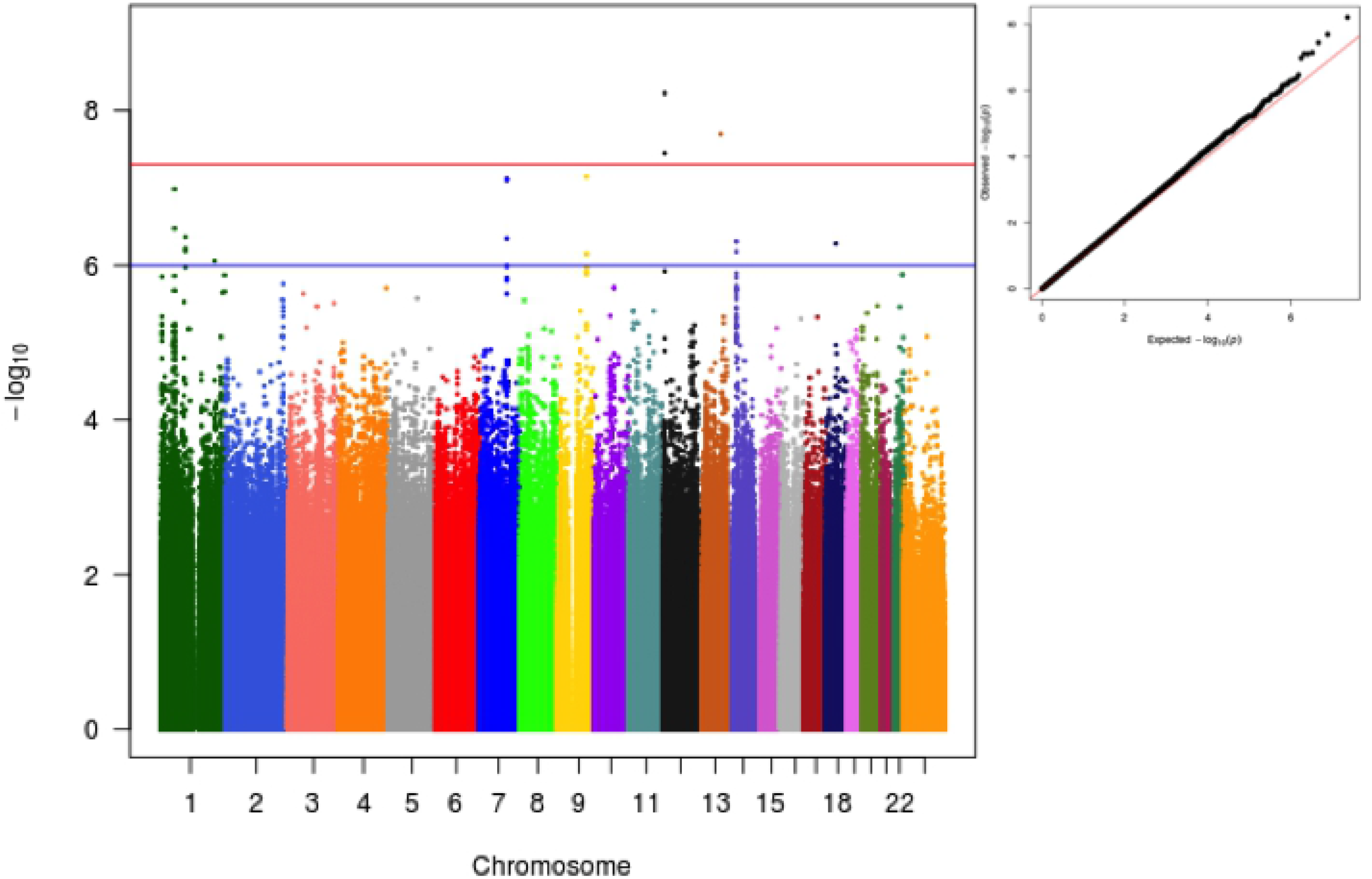
Manhattan and qqplot of GWA analysis for log-transformed PCSK9 as a function of variant, age, sex and the first 4 PCs.

#### Genome-wide significant loci

We observed two genome-wide significant loci and seven loci that reached the pre-defined p value threshold of 1 x 10^−6^ (Table 2).

**Table 2.** Genome-wide and suggestive associated loci.

| Marker | Chromosome | Position | Allele.A | Allele.B | p. value |
| --- | --- | --- | --- | --- | --- |
| chr1:52031405:G:T | 1 | 52497077 | T | G | 1.03E-07 |
| chr1:95303819:A:G | 1 | 95769375 | G | A | 6.11E-07 |
| chr1:206737858:C:G | 1 | 206911203 | G | C | 8.80E-07 |
| chr7:105692603:C:T | 7 | 105333050 | T | C | 4.49E-07 |
| chr9:110739092:T:G | 9 | 113501372 | G | T | 7.18E-08 |
| chr12:4999256:T:C | 12 | 5108422 | C | T | 3.54E-08 |
| chr13:89535768:T:G | 13 | 90188022 | G | T | 2.00E-08 |
| chr14:35161356:C:T | 14 | 35630562 | T | C | 6.64E-07 |
| chr18:37210453:T:C | 18 | 34790416 | C | T | 5.24E-07 |

The strongest associations with PCSK9 plasma levels were observed on chromosome 12p13.32, top lead SNP rs116367042 (P-value 5.97e-09). A regional association plot of the locus is shown in Fig 2. The closest gene *is KCNA5* and significant eQTLs have been observed in the region for *AKAP3*, *DYRK4*, *KCNA5*, *KCNA1*, *NDUFA9*, and *GALTN8*. The region has been described as associated with serum uric acid levels in a previous GWAS.

**Fig 2.**
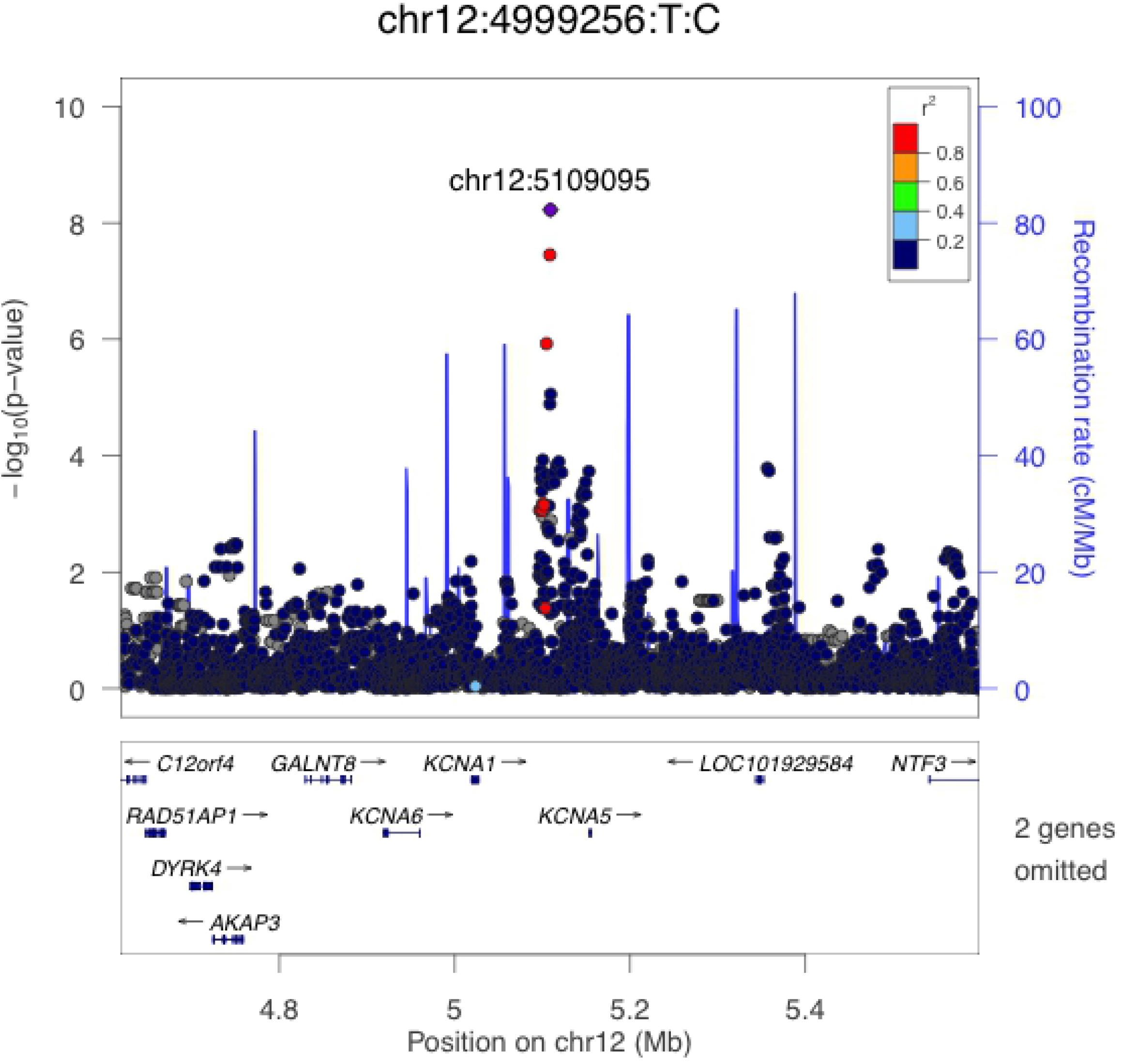
Local association plot for rs116367042.

The second genome-wide significant hit was observed in 13q31.2, in the region coding for *LINC00353*. A regional association plot of the locus is shown in S1A Fig. Notably, only a single marker was associated with PCSK9 levels at this locus, reducing, thus, its credibility.

#### Suggestive loci

Using a pre-defined suggestive significance threshold of 1×10^−6^ we identified additional 7 loci (S2 Table and S2 Fig.). Those loci were on chr1p32.3 (nearest gene *TXNDC12*, SupFig. 2B), chr1p21.3 (nearest gene *RWDD3*, S2C Fig.), chr1q32.1 (nearest gene *MAPKAPK2*, S2D Fig.), chr7q22.3 (nearest gene *ATXN7L1*, S2E Fig.), chr9q31.3 (nearest gene *MUSK*, S2F Fig.), chr14q13.2 (nearest gene *KIAA0391*, S2G Fig.), and chr18q12.2 (nearest gene *KIAA1328*. S2H Fig.). Additional annotations of suggestive loci are available in Supporting information.

### PCSK9 locus association structure

Previous GWAS and candidate-gene association studies have observed significant associations between PCSK9, LDL-C, and TC levels and genetic variants at the PCSK9 locus. Here we extend these observations using a multi-ethnic sample (S3 Fig.). Of note, stronger associations are located at the 3’ region of PCSK9 and within the nearby USP24 gene. Interestingly, previous studies in Europeans, African and other admixed samples have also described stronger associations for total cholesterol and LDL levels at this same region.

Linkage disequilibrium of the PCSK9 locus was resolved in four main haplotype blocks (Supporting information). Fifty-seven markers were nominally associated with PCSK9 levels being the most associated rs505151, rs662145, rs487230, and rs555687 (Supporting information). Tagging associated SNPs in the PCSK9 locus we were able to reduce the number of associated variants from 57 to 20, capturing 100% of the initial variation.

Using information from all 20 tagged markers and a stepwise regression approach we were able to derive a PCSK9 instrumental variable made of four independently associated markers at the PCSK9 locus (cis-pQTLs) (Supporting information). The R-squared for the multiple regression model containing all 4 markers was 0.036. Of particular importance, a model containing independently associated markers, BMI, age and smoking status, although highly significant (*p* = 5.186e-08) was only able to explain 5.2% of the overall variation in PCSK9 plasma levels in our sample, being genetic information the variable with the highest effect size in our model.

### Colocalization analysis of associated loci

Finally, we have studied the colocalization pattern between the identified loci and expression traits of the genes in the vicinity of the association signal. For this we used data from all the available tissues in the GTEx database. Colocalization analysis was able to suggest that *RWDD3*, *ATXN7L1*, *KCNA1*, and *FAM177A1* are potential candidates to be the mediators of the observed associations on chromosome 1, 7, 12 and 14, respectively (S1 Table).

## Discussion

PCSK9 is a serine protease involved in a protein-protein interaction with the LDL receptor that has both human genetic and clinical validation [12]. PCSK9 binds to the LDL-R and is thought to reduce the recycling of these proteins from the cell surface (sending them to lysosomes instead), inhibiting LDL-particle removal from the extracellular fluid [13]. Blocking PCSK9 can lower blood LDL-C concentrations, and low PCSK9 levels are associated with lower LDL-C levels and reduced incidence of atherosclerotic cardiovascular disease. Despite the elusive importance of PCSK9 in lipoprotein homeostasis, few studies have analyzed PCSK9 plasma levels as a function of global genetic variation [14, 15]. Understanding the genetic architecture that modulates PCSK9 levels may help dissect the mechanisms by which PCSK9 inhibition improves vascular function and overall cardiovascular morbidity and mortality.

It is assumed that PCSK9 modulates cardiovascular risk through cholesterol levels, more specifically LDL-C levels. Indeed, pharmacological inhibition of PCSK9 leads to significant decreases in LDL-C and reduction in the incidence of cardiovascular events. However, it is not known whether PCSK9 has other actions independent of plasmatic LDL-C levels [16]. Indeed, PCSK9 is substantially expressed in arterial walls and macrophages [17]. It has also been shown to be associated with metabolic factors other than lipoproteins. It is positively associated with albumin, liver enzymes (ALT, ALP, AST, GGT) and with hepatic steatosis, although whether this association is confounded by or mediated by LDL-C is still unclear [18].

Another important point to be considered is that there is great interindividual variation in both PCSK9 levels and response to PCSK9 inhibitors [8, 19, 20]. In addition, only about 20% of circulating PCSK9 variance can be explained by clinical variables. Previously identified genetic variation only add less than 5% to this figure, almost all of it from eQTL and pQTL within the PCSK9 locus itself [21, 22].

Here we have conducted a GWAS study aiming at identifying genetic determinants of PCSK9 plasma levels. To our knowledge this is the second GWAS conducted for PCSK9 levels and the first using a sample from a multi-ethnic population. Despite the relatively small sample size we were able to observe two genome-wide significant association loci and a number of loci with suggestive association signals. In addition, we have confirmed the previously described association between PCSK9 levels and common genetic variation at the *PCSK9* locus [14].

We have not been the first to describe genome-wide significant variants associated with plasma PCSK9 levels outside of the PCSK9 gene. Pott et al. in a GWAS conducted in 3290 individuals from the LIFE-Heart cohort identified variations within the *FBXL18* gene to be associated with PCSK9 levels [14]. We did not identify any association in this region and together with the low imputation quality the authors of this previous GWAS described, we suggest the association between *FBXL18* and PCSK9 to be a false-positive signal.

Plasma PCSK9 levels have been associated with several cardiovascular and metabolic risk factors [23]. Notwithstanding the understanding of PCSK9 mechanism at the molecular level, completely understanding the directionality of the associations between PCSK9 levels and other metabolic traits has been ill explored. In fact, most studies assume that PCSK9 associated with lipid levels because of the interaction between PCSK9 and LDLR at the molecular level. However, it is unknown whether different predictors of PCSK9 levels are indeed associated with the same degree of increased cardiovascular risk. Indeed, recent experimental and clinical studies have also reported that higher circulating PCSK9 levels contributed to coronary atherosclerosis by enhancing the expression of pro-inflammatory genes, promoting apoptosis of human endothelial cells and activating platelet reactivity [24, 25]. The causal directionality of these associations however, has not been fully explored.

By identifying potentially trans associations with PCSK9 levels, our data give rise to the possibility of a more complex mechanism, where different genetic factors may modulate PCSK9 levels. It remains to be determined if PCSK9 levels driven by trans genetic factors carry the same increased risk of cardiovascular disease as PCSK9 levels determined by genetic variation at the *PCSK9* locus. In summary, our data suggest that PCSK9 levels may be modulated by upstream targets other than genetic variation in the PCSK9 gene, which are well-known proxies for PCSK9 levels.

This study has some potential limitations. First and foremost, we have not been able to find a suitable replication sample for our GWAS results. The observed genome-wide significant loci still need to be replicated in an independent sample to be, in effect, taken as drivers of PCSK9 plasma levels. In addition, the reduced sample size of our study may have prevented us to identify other genome-wide significant loci with decreased effect size. Mendelian randomization analysis using our sample lacked the necessary statistical power to derive robust conclusions regarding the causality of trans PCSK9 variants and coronary artery disease or even LDL-C levels. These aspects should be better defined in further studies.

In conclusion, we describe new genome-wide significant loci associated with PCSK9 plasma levels in a sample from a healthy population. Our results suggest that PCSK9 levels may be modulated by trans genetic variation outside of the *PCSK9* gene. Understanding both environmental and genetic predictors of PCSK9 levels may help in identifying new targets for cardiovascular disease treatment and contribute to better assessment of the benefits of long-term PCSK9 inhibition treatment.

## Supporting Information

**S1 Fig. Log-PCSK9 GWA adjusted for age.**

**S2 Fig. Suggestive associated loci.** Those loci were on 13q31.2, in the region coding for *LINC00353* (A); chr1p32.3 (nearest gene *TXNDC12*, B), chr1p21.3 (nearest gene *RWDD3*, C), chr1q32.1 (nearest gene *MAPKAPK2*, D), chr7q22.3 (nearest gene *ATXN7L1*, E), chr9q31.3 (nearest gene *MUSK*, F), chr14q13.2 (nearest gene *KIAA0391*, G), and chr18q12.2 (nearest gene *KIAA1328*, H).

**S3 Fig. Local association structure at the PCSK9 gene locus (upper panel); local linkage-disequilibrium structure at the PCSK9 gene locus.**

**S1 Table. Multiple linear regression for PCSK9 instrumental variable.**

**S2 Table. Colocalization analysis with GTEX.**

